# Robust discovery of mutational signatures using power posteriors

**DOI:** 10.1101/2024.10.23.619958

**Authors:** Catherine Xue, Jeffrey W. Miller, Scott L. Carter, Jonathan H. Huggins

**Affiliations:** Harvard University, Department of Biostatistics; Dana Farber Cancer Institute, Department of Data Science; Boston University, Department of Mathematics & Statistics

## Abstract

Mutational processes, such as the molecular effects of carcinogenic agents or defective DNA repair mechanisms, are known to produce different mutation types with characteristic frequency profiles, referred to as mutational signatures. Non-negative matrix factorization (NMF) has successfully been used to discover many mutational signatures, yielding novel insights into cancer etiology and targeted therapies. However, the NMF model is only a rough approximation to reality, and even small departures from this assumed model can have large negative effects on the accuracy and reliability of the results. We propose a new approach to mutational signatures analysis that improves robustness to misspecification by using a power posterior for a fully Bayesian NMF model, while employing a sparsity-inducing prior to automatically infer the number of active signatures. In extensive simulation studies, we find that our proposed approach recovers more true signatures with greater accuracy than current leading methods. On whole-genome sequencing data for six cancer types from the ICGC/TCGA Pan-Cancer Analysis of Whole Genomes Consortium, we find that our method is able to accurately recover more signatures than the current state-of-the-art.

## 1. Introduction

Carcinogenic processes such as UV radiation, smoking, defective DNA repair mechanisms, and naturally occuring biochemical reactions generate characteristic patterns of somatic mutations known as mutational signatures (Helleday et al., 2014; Koh et al., 2021; Nik-Zainal et al., 2012b). While these processes cannot be observed directly in patients, the cumulative effect of multiple processes on an individual tumor can be quantified using genome sequencing, and their distinct mutational signatures can be inferred using statistical modeling. Non-negative matrix factorization (NMF) models have proven effective in estimating mutational signatures as well as the mutational load due to each signature in each tumor sample (Alexandrov et al., 2013b; Nik-Zainal et al., 2012a). Mutational signatures analysis has contributed to novel insights in a variety of areas of cancer research (Alexandrov et al., 2013a, 2015, 2020; Li et al., 2020; Nik-Zainal et al., 2012b) and has emerging translations to clinical outcomes (Chakravarty and Solit, 2021).

Existing methods for mutational signatures analysis are fundamentally limited by the fact that they assume—either explicitly or implicitly—a particular probabilistic model for how mutations arise. However, any assumed model will only be a rough approximation to reality. Unfortunately, using an incorrect model—known as *model misspecification*—can lead to spurious inferences (Miller and Dunson, 2018). In particular, a key challenge in mutational signature discovery is determining the number of active signatures (Alexandrov et al., 2013a,b; Kim et al., 2016; Rosales et al., 2017), and standard statistical methods for this type of model selection problem tend to be extremely sensitive to misspecification (Cai et al., 2021; Miller and Dunson, 2018). For instance, methods based on automatic relevance determination (ARD) (Kim et al., 2016; Tan and Févotte, 2013) and Bayesian spike-and-slab models (Legramanti et al., 2020) tend to overestimate the number of signatures when there is mild overdispersion (Zito and Miller, 2024). Of course, robustness to outliers can be obtained using overdispersed models such as negative binomial (Lyu et al., 2020), but this does not handle any other types of misspecification. Perhaps due to this sensitivity, the current leading methods rely heavily on *ad hoc* techniques such as manual filtering (Alexandrov et al., 2020) or neural networks (Nebgen et al., 2021) for determining the number of signatures.

In this paper, we show that leading methods for mutational signatures analysis are not robust to model misspecification, and we introduce a novel method that exhibits better performance under misspecification. Specifically, we investigate the degree to which misspecification causes existing methods to (1) fail to find important processes or (2) infer spurious processes that do not actually exist. Our findings indicate a lack of robustness in two widely used methods: SigProfilerExtractor (Alexandrov et al., 2013b, 2020; Islam et al., 2022) and SignatureAnalyzer (Alexandrov et al., 2020; Kim et al., 2016; Tan and Févotte, 2013); see Figure 1 for examples. To address these limitations, we propose a new method called *BayesPowerNMF*. By leveraging a power posterior (Miller and Dunson, 2018) and a sparsity-inducing prior (Zito and Miller, 2024), BayesPowerNMF provides nonparametric robustness to model misspecification and automated, principled selection of the number of latent processes. Through simulation studies and a comparison to existing methods on whole-genome data for six cancer types (The ICGC/TCGA Pan-Cancer Analysis of Whole Genomes Consortium, 2020), we show that our BayesPowerNMF method finds more true processes and fewer spurious processes than competitors; see Figure 1 for two illustrations.

**Figure 1.**
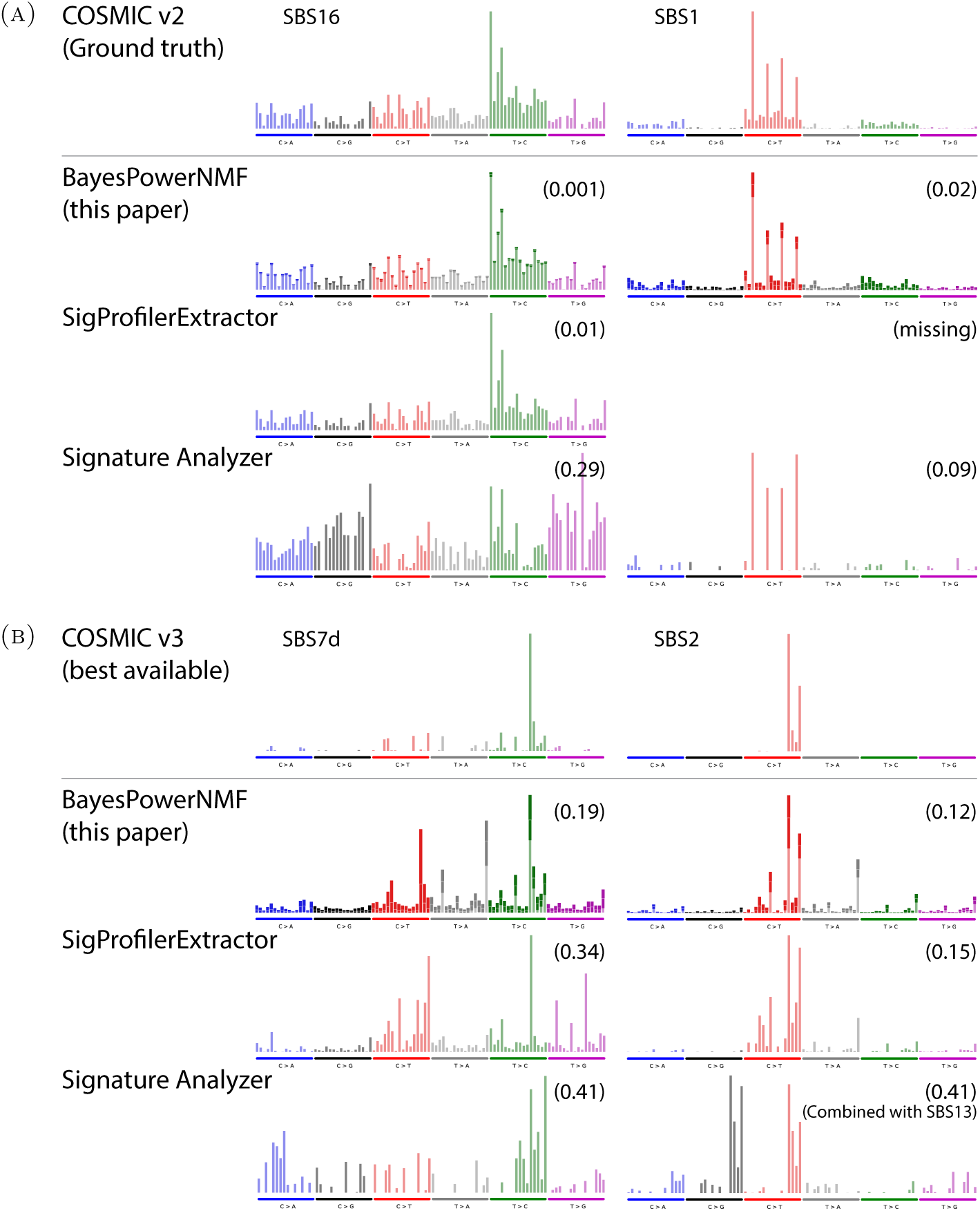
**(A)** Examples from simulated misspecified liver cancer data. For two ground truth signatures used to generate the data (SBS16 and SBS1 from COSMIC v2, shown in the top row), we show the best-matching signatures inferred by two leading methods (SigProfilerExtractor and SignatureAnalyzer) and our proposed method (BayesPowerNMF). The cosine errors between the inferred and true signatures are in parentheses. When the model is misspecified, SigProfilerExtractor tends to miss more signatures (such as SBS1 in this case) and SignatureAnalyzer tends to have significantly higher cosine errors compared to BayesPowerNMF. **(B)** Examples from real melanoma data. Same as in (A), but comparing with the closest match to SBS7d and SBS2 from COSMIC v3. For BayesPowerNMF, the bolded section of each bar indicates a 95% credible interval, quantifying the uncertainty in each signature. Meanwhile, SigProfilerExtractor and SignatureAnalyzer only provide point estimates that may be misleading in cases such as SBS7d for which there is high uncertainty.

The article is organized as follows. In Section 2, we introduce our proposed methodology. We present simulation results comparing our method to leading existing methods in Section 3. Then, in Section 4, we compare the methods in an application to whole-genome data from the Pan-Cancer Analysis of Whole Genomes Consortium (PCAWG). We conclude with a discussion, including limitations and possible directions for future work, in Section 5.

## 2. Methodology

In this section, we describe the standard Poisson NMF model, our sparsity-inducing prior, the power posterior, our technique for choosing the power, and the overall workflow of our BayesPowerNMF method. NMF is based on the assumption that the data matrix ***X*** = (*X*_nm_) ∈ R^N*×*M^ can be approximated by a low-rank factorization of the form

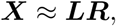

where the (unknown) matrices ***L*** = (*L*_nk_) ∈ R^N*×*K^ and ***R*** = (*R*_km_) ∈ R^K*×*M^ have non-negative entries (Fu et al., 2019; Lee and Seung, 2000). Here, R_+_ = [0, ∞) and we use bold font to indicate matrices. Since arbitrary multiplicative constants can be moved between ***L*** and ***R*** without affecting the product ***LR***, we constrain the rows of ***R*** to sum to 1, that is, *R*_km_ = 1 for all *k* = 1*, …, K*. Low-rank factorizations such as this often provide valuable insights into the latent structure giving rise to the data.

In mutational signatures analysis, the rows of ***X*** correspond to samples *n* = 1*, …, N*, the columns of ***X*** correspond to a pre-defined set of *M* non-overlapping mutation types, and each entry *X*_nm_ ∈ {0, 1, 2*, …*} represents the number of times that mutation type *m* is observed in sample *n*. The idea of the NMF model is that mutations arise due to multiple unknown mutational processes, and each such process generates mutations according to a distinct profile of frequencies. The *k*th row of ***R***, say *R*_k_ = (*R*_k1_*, …, R*_kM_), represents the frequencies with which mutation types *m* = 1*, …, M* occur under the *k*th mutational process, and *R*_k_ is referred to as the *mutational signature* of this process. Thus, *K* represents the number of mutational processes represented in the model, (that is, the number of latent factors). The loading *L*_nk_ represents the activity of process *k* in sample *n*.

### 2.1. Model and sparsity-inducing prior

Our methodology starts with the *Poisson NMF model*

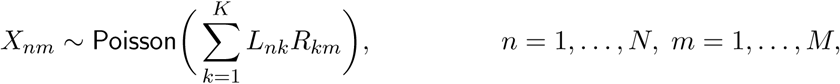

independently; or in more compact matrix notation,

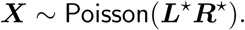

This model is standard in mutational signatures analysis, and it can be justified from first principles (Zito and Miller, 2024). The parameters of the model are (***L****, **R***) and the likelihood is

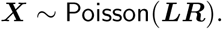

For the prior on ***R***, we use independent *M*-dimensional Dirichlet priors on the rows,

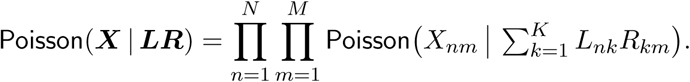

where *α*_0_ *>* 0. For the prior on ***L***, since the number of active mutational processes is not known *a priori*, we employ a sparsity-inducing prior introduced by Zito and Miller (2024). Specifically, we place independent gamma priors on the loadings,

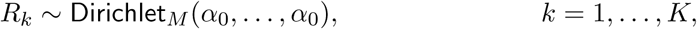

where *a >* 0 is the shape and *a/µ*_k_ is the rate, so that E(*L*_nk_) = *µ*_k_ and hence *µ*_k_ is the mean prior loading for the *k*th mutational process. Finally, we give *µ*_k_ an inverse gamma hyperprior,

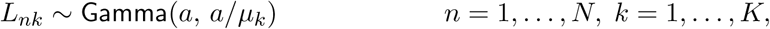

independently, where *a*_0_ = *N*_0_*a* + 1, *b*_0_ = *ε*(*a*_0_ − 1), and *ε >* 0 is a small nonzero constant. With these choices, the prior mean of *µ*_k_ becomes E(*µ*_k_) = *ε* and its full conditional mean is

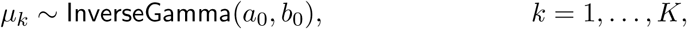

This choice of hyperprior has a compressive property that induces column-wise sparsity in the loadings matrix under the posterior distribution. More precisely, given the data ***X***, it shrinks the *µ*_k_ value for inactive processes down to *µ*_k_ ≈ *ε*, while having a moderate shrinkage effect on the *µ*_k_ value for active processes. Consequently, the loadings *L*_nk_ for inactive processes shrink to small values ≈ *ε*, while the loadings for active processes are only slightly affected by the hyperprior (Zito and Miller, 2024).

In our experiments, we set *α*_0_ = 0.5, *a* = 0.5, *ε* = 0.001, and *N*_0_ = 10. Choosing *α*_0_ = 0.5 and *a* = 0.5 makes the priors on *R* and *L* weakly informative. Specifically, the prior on *R*_k_ is the Jeffreys prior for a multinomial likelihood and the conditional prior variance of *L*_nk_ is Var(*L*_nk_ | *µ*_k_) = 2*µ*^2^. The value of *ε* does not matter much as long as it is significantly smaller than the smallest true loadings. The choice of *N*_0_ = 10 was found to work well empirically, providing sparsity without shrinking the *µ*_k_ values too strongly.

### 2.2. Power posterior

The standard Poisson NMF model works well on data generated from the model itself, but its performance suffers when the model is not exactly correct, as we demonstrate in Section 3. To address this issue, we employ the power posterior technique for improving robustness to misspecification (Miller and Dunson, 2018). Letting *π*_0_(*θ*) denote the prior density on *θ* = (***L****, **R**, µ*), the standard Bayesian posterior is

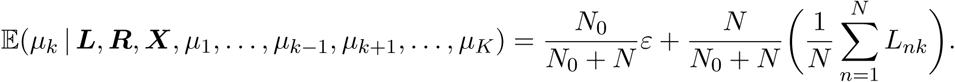

where *p*(***X*** | *θ*) = Poisson(***X*** | ***LR***) and *Z*(***X***) = *p*(***X*** | *θ*) *π*_0_(*θ*) d*θ*. For *ξ* ∈ [0, 1], the *power posterior* is defined as

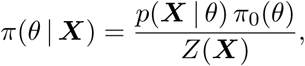

where *Z*_ξ_(***X***) = *p*(***X*** | *θ*)^ξ^ *π*_0_(*θ*) d*θ* is the normalization constant. In particular, when *ξ* = 0 or *ξ* = 1, we recover the prior or the standard posterior, respectively; that is, *π*_0_(*θ* | ***X***) = *π*_0_(*θ*) and *π*_1_(*θ* | ***X***) = *π*(*θ* | ***X***). As shown by Miller and Dunson (2018), using a power posterior with 0 *< ξ <* 1 provides nonparametric robustness for misspecified latent variable models – particularly when the latent dimensionality is unknown, such as in mutational signatures analysis. Furthermore, Medina et al. (2022) showed that—compared to using the standard posterior—using the power posterior of a misspecified model is closer to the standard posterior of a more flexible well-specified model, in terms of Kullback–Leibler divergence.

To perform inference, we use the Stan probabilistic programming system (Carpenter et al., 2017) to draw Markov chain Monte Carlo (MCMC) samples from the power posterior of the NMF model. Specifically, we define the log target density to be

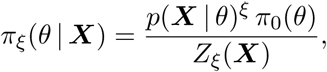

Using Stan makes it a simple matter of multiplying the log-likelihood part of this function by *ξ* in order to target the power posterior rather than the standard posterior. For the MCMC sampler settings, we use four chains, each with 10,000 samples after a burn-in of 10,000 iterations. We choose the best of these four chains, defined as the one with the largest approximate marginal likelihood, using the approximation of Pritchard et al. (2000).

### 2.3. Choosing the power

The power *ξ* should be selected to reflect the severity of the model misspecification (Medina et al., 2022; Miller and Dunson, 2018). When there is no misspecification—that is, when the data are generated by the assumed model—we would ideally take *ξ* = 1, which corresponds to the standard Bayesian posterior. When there is misspecification, smaller values of *ξ* are preferable. However, it can be difficult to choose *ξ* in a data-driven way.

We propose the following simulation-driven approach to choosing *ξ*. First, select a set of *pilot signatures* (for example, we use a relevant subset of the COSMIC v2 signatures^1^; Alexandrov et al., 2015; Nik-Zainal et al., 2016) and estimate loadings to fit the input data using these pilot signatures. Next, generate pilot data from several plausible data generating processes (DGPs) based on the pilot signatures and corresponding estimated loadings. Then, fit each pilot data set using a range of *ξ* values, and select a value of *ξ* that performs well across all of the pilot data sets.

By exploring simulated data from DGPs that are not equal to the assumed model, but are anchored at parameters fit to the input data, we are able to select a *ξ* that yields robustness to plausible perturbations within an appropriate data-driven neighborhood of the model. This yields robustness to any DGPs that would require a similar power *ξ*, not just to the specific forms of DGP used to generate pilot data. Thus, it is not critical to get the DGPs exactly right – just to make them sufficiently rich and plausible.

### 2.4. Simulation data generating processes

Given a matrix of pilot signatures ***R***^*^ = (*R*^*^) ∈ R^K*×*M^ and corresponding matrix of loadings ***L***^*^ = (*L*^*^) ∈ R^N*×*K^, we generate pilot data sets using the following four DGPs based on ***R***^*^ and ***L***^*^:

1. The well-specified DGP generates data from the assumed Poisson NMF model, as in Equation (1):

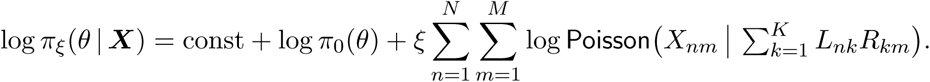
2. The *α*-contaminated DGP generates data from the Poisson NMF model, but with an additional “contamination signature” *R*^-^_n_: for *α* ∈ (0, 1),

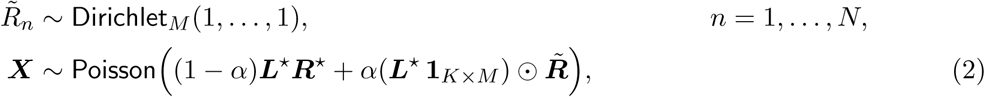

where **1**_K*×*M_ denotes the *K* × *M* matrix of ones, ⊙ denotes entry-wise multiplication, and ***R***^-^ is the matrix with rows *R*^-^_1_*, …, R*^-^_N_. The contamination signature *R*^-^_n_ is different for each sample, to reflect the possibility of tumor-specific variability arising from microenvironment effects or rare mutational processes. Equation (2) gives *R*^-^_n_ a loading of *α*, and the existing loadings *L*^*^ are downweighted by a factor of 1 − *α* to keep the total loading invariant.
3. The *γ*-perturbed DGP generates each sample using slightly perturbed versions of each signature, with the perturbations being sample-specific: given *γ >* 0 and *β*_1_*, …, β*_K_ *>* 0,

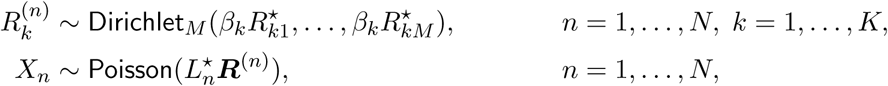

where *L*^*^ is the *n*th row of ***L***^*^, ***R***^(n)^ is the matrix with rows *R*^(n)^*, …, R*^(n)^, and *X*_n_ is the *n*th row of ***X***. The constants *β*_k_ control the size of perturbations, and are set such that the expected *cosine error* between *R*^(n)^_k_ and *R*^*^_k_ is approximately *γ*, where the cosine error is defined as 1−*R*^T^_k_*R*^*^_l_*/*(∥*R*_k_∥∥*R*^*^_l_∥); see Appendix B for details. This DGP reflects the possibility that each mutational process might behave slightly differently across tumors, for instance due to the tumor microenvironment, different failure modes of the same DNA repair mechanism, or smoking different brands of cigarette.
4. The *κ*-overdispersed DGP generates data with higher variance than the well-specified data, to reflect the fact that there is often additional variability due to sample- and mutation-type specific effects that may be biological or technical: given *κ >* 1,

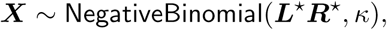

where NegativeBinomial(*µ, κ*) denotes the negative binomial distribution with mean *µ* and variance *κµ*.

In our experiments, we use *α* = 0.02, *γ* = 0.0025, and *κ* = 2 for the settings of these DGPs. These settings are chosen to reflect plausible levels of biological and technical perturbations from the Poisson model. Specifically, *α* = 0.02 equates to 2% of observed mutations originated from some source of contamination and *γ* = 0.0025 is a threshold where the simulated signatures are almost always still recognizable as coming from the original reference signature (99% of simulated signatures have cosine error *<* 0.1). We set *κ* = 2 to reflect the empirical overdispersion seen in loadings through exploratory analysis of real count data.

### 2.5. Overall workflow

To summarize, the overall workflow of our proposed method, *BayesPowerNMF*, consists of the following five stages (which are depicted in Figure 2):

1. *Define pilot signatures and loadings.* To tailor the pilot data sets to the input data set, we define the pilot signatures ***R***^*^ and loadings ***L***^*^ as follows. Using non-negative least squares (Lawson and Hanson, 1987), we estimate loadings for the 30 COSMIC v2 reference signatures (Alexandrov et al., 2015; Nik-Zainal et al., 2016) for each individual in the cohort. We then drop any reference signatures that, based on the estimated loadings, do not contribute at least 2% of the total mutation count in the cohort or at least 10% of the mutation count of any individual, so that each remaining signature is well represented. The remaining loadings are not re-fit, to maintain the accuracy of the loadings.
2. *Generate pilot data sets.* For each of four DGPs, generate a pilot data set(s) based on parameters ***L***^*^ and ***R***^*^, which results in *J* total simulated data sets ***X***_1_*, …, **X***_J_. As described in Section 2.4, we use *J* = 4: the well-specified DGP plus the three misspecified DGPs with parameters *α* = 0.02, *γ* = 0.0025, and *κ* = 2.
3. *Estimate parameters for a range of ξ values.* For each power *ξ* in a range of candidate values, for each pilot data set, we estimate ***L*** and ***R*** using the mean of the power posterior (Section 2.2) and match the estimated signatures *R*_k_ to the pilot signatures *R*^*^. Matching *R*_k_ to *R*^*^ is done using the Hungarian algorithm for optimal bipartite matching (Kuhn, 1955) with cosine error as the assignment cost.
4. *Choose ξ to maximize accuracy across pilot data sets.* We choose the largest power *ξ* that accurately recovers ***L***^*^ and ***R***^*^ according to the following metrics (see Figure S4): *(i)* recovering as many true signatures as possible with cosine error *<* 0.3, and *(ii)* not inferring any *spurious signatures* that fail to match any true signature.
5. *Apply power posterior to the input data.* Using the selected power *ξ*, we apply the power posterior to the original input data ***X*** to estimate ***L*** and ***R***. This yields the final estimates of ***L*** and ***R*** produced by the workflow.

**Figure 2.**
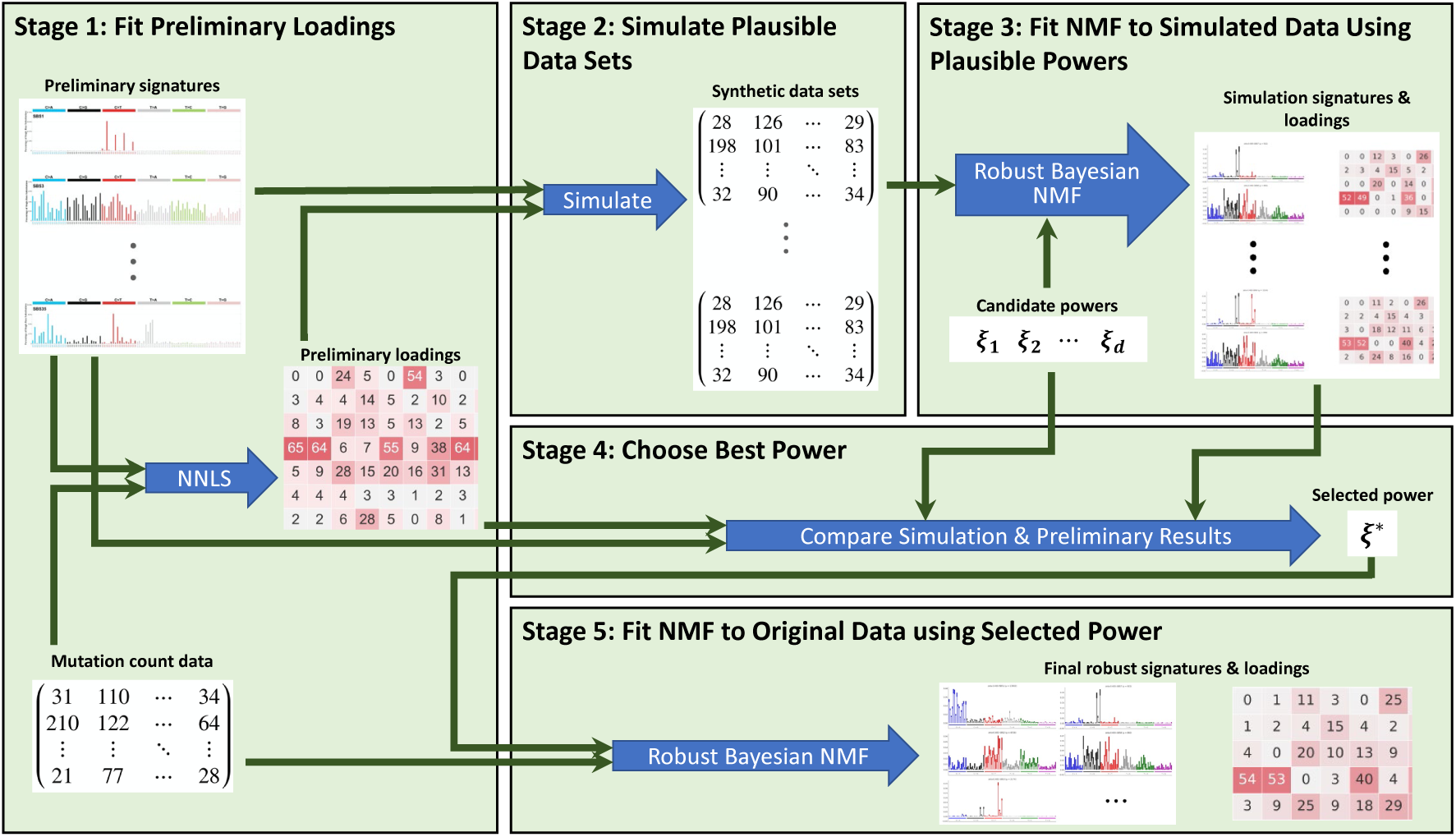
BayesPowerNMF workflow for robust mutational signature discovery. **(1)** Pilot signatures from a reference (such as COSMIC) are used to estimate loadings to fit the input data. **(2)** These signatures and loadings are used to simulate pilot data sets from DGPs representing plausible perturbations of the model. **(3)** The power posterior for the Bayesian NMF model is applied to each combination of candidate power *ξ*_i_ and simulated data set ***X***_j_. **(4)** We identify the power that best recovers the pilot signatures and loadings across all of the pilot data sets. **(5)** Using the selected power, we apply the power posterior for the Bayesian NMF model to the original input data.

## 3. Simulations

In this section, we compare performance on synthetic data simulating six cancer types, using the four data generating processes described in Section 2.4. We compare three NMF methods for inferring mutational signatures: (1) SigProfilerExtractor, a version of the algorithm used to define the COSMIC signatures (Alexandrov et al., 2013b; Islam et al., 2022), (2) SignatureAnalyzer, based on a Bayesian point estimation algorithm (Tan and Févotte, 2013) used in several previous publications (Haradhvala et al., 2018; Kasar et al., 2015; Kim et al., 2016), and (3) BayesPowerNMF, our proposed method.

### 3.1. Simulated data

We generate synthetic data sets in the well-specified case for six cancer types (lung adenocarcinoma, stomach, melanoma, ovary, breast, and liver) and in the three misspecified cases (*α*-contaminated, *γ*-perturbed, and *κ*-overdispersed) for three cancer types (lung adenocarcinoma, ovary, and liver). These simulated data sets are generated as in stages 1 and 2 of the BayesPowerNMF workflow in Section 2.5, with the input data set being a mutation count matrix for the corresponding cancer type from the PCAWG project (Alexandrov et al., 2020). In addition, we consider randomly subsampled data sets of sizes 20, 30, 50, 80, 120, 170, and 230 from the simulated well-specified liver data set of 326 samples. Thus, altogether, starting from 6 real data sets (one for each cancer type), we generate 15 complete synthetic data sets (6 well-specified and 3 × 3 misspecified) plus 104 data sets subsampled from the complete well-specified liver data set (32 of size *N* = 20, 20 of size *N* = 30, 12 of size *N* = 50, and 10 each of the remaining sizes). Using each of these synthetic data sets as input, we then apply each of the three competing methods listed above: SigProfilerExtractor, SignatureAnalyzer, and BayesPowerNMF (following our workflow in Section 2.5).

### 3.2. Performance evaluations

We evaluate performance by measuring how well the groundtruth signatures and loadings are recovered in terms of several metrics. Here, ground truth is known since the data is simulated. First, we compute the optimal matching (in terms of cosine error) between estimated and true signatures. We then quantify how accurately each signature is recovered by computing the cosine error between each pair of matched signatures. We compute the precision and recall for recovering the true signatures by setting a threshold on cosine error and considering a true signature to be correctly recovered if the cosine error of its matching estimated signature is within this threshold. Precision is defined as (# correctly recovered) */* (# estimated signatures) and recall is (# correctly recovered) */* (# true signatures). Plots of precision and recall are then generated by varying the threshold. Additionally, we use bubble plots to display all of the matched pairs of signatures, along with their cosine errors and loadings, for each method and each data set. This provides a convenient visual summary of the results in a single plot.

### 3.3. Simulation results

#### 3.3.1. Overall precision and recall

Figure 3 shows the precision and recall for each of the three methods, over a range of cosine error thresholds from 0 to 0.2. These curves are averaged over all of the well-specified data sets (left side) and misspecified data sets (right side). Our BayesPowerNMF method provides the highest precision and recall at nearly every threshold. In particular, BayesPowerNMF has much higher recall in the misspecified settings. SigProfilerExtractor performs only somewhat worse than BayesPowerNMF in the well-specified settings, but its recall suffers under misspecification.

**Figure 3.**
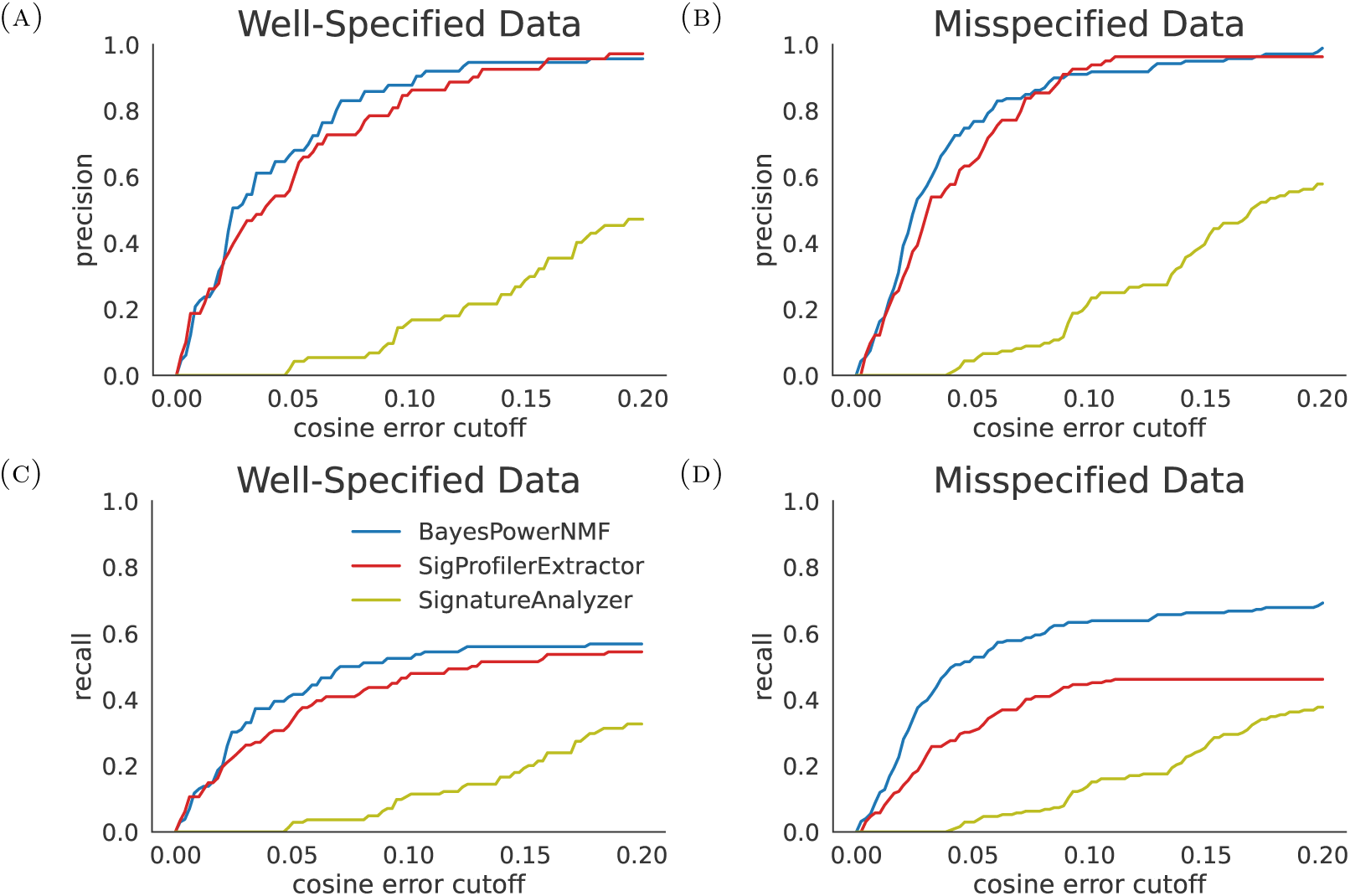
(Top) Precision curves for each method as a function of cosine error threshold, averaged across the simulated well-specified data sets (left) and misspecified data sets (right). (Bottom) Same for the recall curves. BayesPowerNMF has the highest precision and recall overall. SigProfilerExtractor performs nearly as well as BayesPowerNMF in the well-specified settings, but its recall is lower under misspecification. SignatureAnalyzer exhibits low precision and recall in these simulations.

The signatures missed by SigProfilerExtractor tend to be ones with smaller true loading; see Figure S2. One reason for this may be because SigProfilerExtractor employs a consensus bootstrap approach that requires agreement across the estimates based on different bootstrap data sets. To compare the performance for each signature individually, we plot the cosine error from the ground truth for SigProfilerExtractor versus BayesPowerNMF in Figure S3; this illustrates that BayesPowerNMF tends to exhibit better performance on a signature-by-signature basis.

We find that SignatureAnalyzer exhibits worse performance across all scenarios, both in terms of precision and recall. The reason for this is that SignatureAnalyzer estimates many signatures, but the estimated signatures have a high cosine error relative to the matching true signatures. Although SignatureAnalyzer is based on a Bayesian model, it employs *maximum a posteriori* estimation rather than posterior uncertainty quantification, and does not benefit from the robustness properties of the power posterior.

#### 3.3.2. Lung adenocarcinoma results

To understand what each method is doing at a more granular level, Figure 4 shows the cosine error and loading for each signature. The true COSMIC signatures are listed on the left, and each column summarizes the results from a given method on a given data set. The size of each bubble represents the loading given by that method to the estimated signature that was matched to the true signature listed on the left. The shade of the bubble represents the cosine error between the estimated signature and the matching true signature. The left-most column (labelled GT) shows the ground truth loadings used to generate the simulated data, and we order the rows (signatures) according to these ground truth loadings. Therefore, a method is performing well on a given data set if *(i)* it has bubbles for the same signatures as GT, *(ii)* the sizes of the bubbles are similar to GT, and *(iii)* the shade of the bubble is white or light gray. Methods with bubbles above the red line estimated a spurious signature that does not match to any of the true signatures used to generate the data.

**Figure 4.**
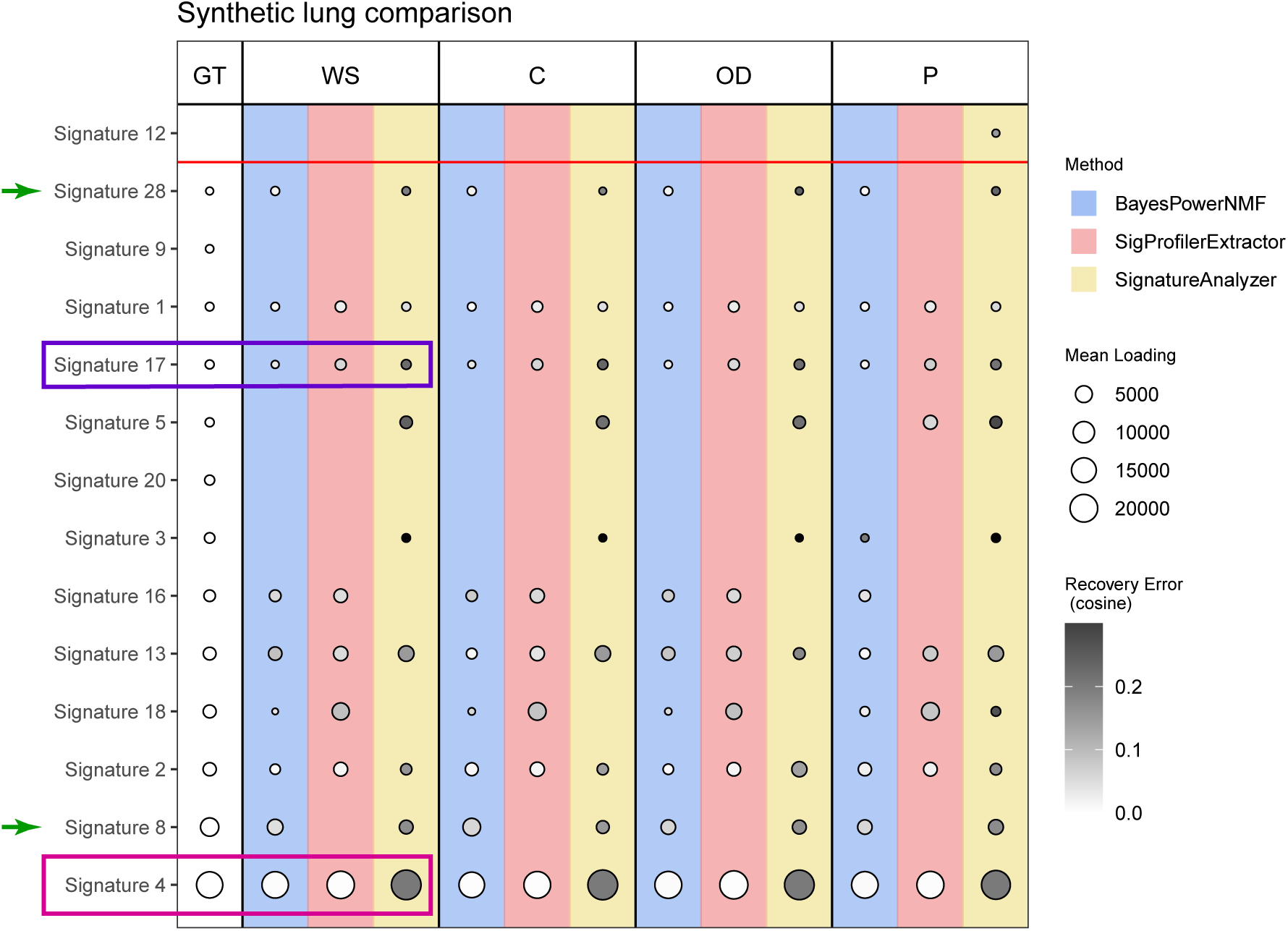
Comparison of results on simulated lung adenocarcinoma data sets, via a bubble plot showing the loading and cosine error of the estimated signatures. Rows represent the true COSMIC v2 signatures, ordered by the ground truth (GT) loading. Each column shows the results for a given method (blue = BayesPowerNMF, red = SigProfilerExtractor, yellow = SignatureAnalyzer) on a given data set (WS = well-specified, C = *α*-contaminated, OD = *κ*-overdispersed, P = *γ*-perturbed). Presence of a bubble means an estimated signature matched that true signature. Bubble size = estimated loading, bubble shade = cosine error between estimated and true, and a bubble above the red line represents an estimated signature that did not match any true signature used to generate the data. Overall, BayesPowerNMF consistently finds the most signatures and has the lowest cosine errors.

For the simulated lung adenocarcinoma data sets, which have *N* = 38 samples, Figure 4 shows that BayesPowerNMF tends to recover more true signatures with lower cosine error than the other methods. SigProfilerExtractor performs second best, but misses some signatures (in particular, Signatures 8 and 28, indicated by green arrows) and tends to have slightly higher cosine error. Note that Signature 8 was not recovered by SigProfilerExtractor in any of the cases, even though this signature has a large true loading (second largest out of all the signatures). SignatureAnalyzer does recover many signatures that match to the true signatures, but they have very high cosine error, even for signatures with high true loading such as Signature 4. On the *γ*-perturbed data, SignatureAnalyzer also produces a spurious signature that matches to COSMIC Signature 12, which was not used to generate the data.

Figure 5a shows true Signature 17 and the matching estimated signatures for each method on the well-specified (WS) data (corresponding to the purple box in Figure 4). This is an example of a signature with relatively small true mean loading and, in comparison with Signature 4, it also has many more near-zero values in the signature itself. BayesPowerNMF recovers the signature with the lowest cosine error (0.03), followed by SigProfilerExtractor (0.06), and then SignatureAnalyzer (0.22). BayesPowerNMF also provides uncertainty quantification in the signature vector, indicated in the plot by the darker region at the top of each bar, which represents a 95% credible interval.

**Figure 5.**
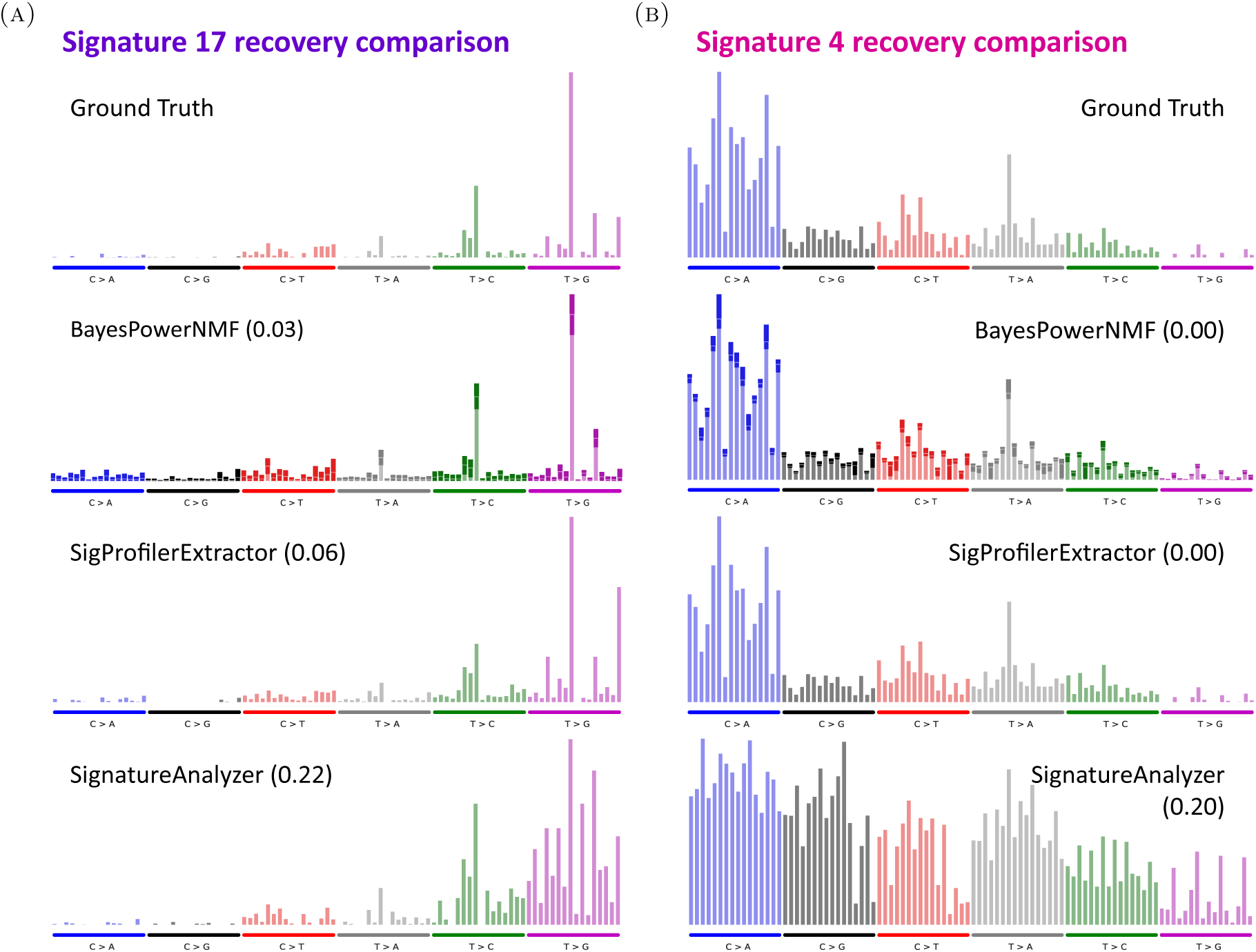
Comparison of estimated signatures on simulated lung adenocarcinoma data sets. **(A)** True Signature 17 and matching estimated signatures (along with their cosine error, in parentheses) for each method on the WS data. For BayesPowerNMF, the bolded section of each bar for represents 95% credible interval and the thin white line in the middle represents the posterior mean. **(B)** Same panel A but for Signature 4.

Similarly, Figure 5b shows true Signature 4 and the matching estimated signatures on the WS data (corresponding to the magenta box in Figure 4). This is an example of a signature with large true mean loading (the largest in this simulated data). All three methods estimate a signature that matches to Signature 4, with estimated loading similar to the true loading.

Furthermore, BayesPowerNMF and SigProfilerExtractor both recover the true signature with very low cosine error (*<* 0.005). However, SignatureAnalyzer’s matching signature has a high cosine error (0.2). For BayesPowerNMF, the 95% credible regions (bolded regions of the bars) tend to indicate less uncertainty in the estimates for Signature 4 compared to 17, especially in the mutation types with smaller rates; this makes sense since Signature 4 has higher true mean loading and thus there is more information about it in the data.

#### 3.3.3. Subsampled liver cancer data sets of varying size

Next, we evaluate the effects of sample size and sampling variability by considering the subsampled data sets of varying size taken from the complete simulated well-specified liver cancer data set of *N* = 326 samples. The full cohort of *N* = 326 samples was generated using 21 ground truth signatures. From this, we repeatedly subsampled data sets of various sizes and ran each method on each subsampled data set. Figure 6 summarizes the results for BayesPowerNMF and SigProfilerExtractor on these data; SignatureAnalyzer is excluded from this analyses due to its overall poor performance. Figure 6a plots the precision and recall curves as a function of cosine error threshold, like in Figure 3. As before, we see that BayesPowerNMF outperforms SigProfilerExtractor, particularly in terms of recall. SigProfilerExtractor does exhibit slightly higher precision at higher cosine error thresholds, but this comes at the expense of much lower recall. Figure 6b shows the number of signatures inferred by each method as a function of the size of the data set. This shows that BayesPowerNMF requires a much smaller sample size to recover the same number of signatures as SigProfilerExtractor. For instance, BayesPowerNMF recovers an average of 9 signatures on a data set of size *N* = 30, whereas SigProfilerExtractor requires *N* = 230 to recover this many signatures on average. Thus, BayesPowerNMF accurately recovers the same number of signatures with a fraction of the sample size.

**Figure 6.**
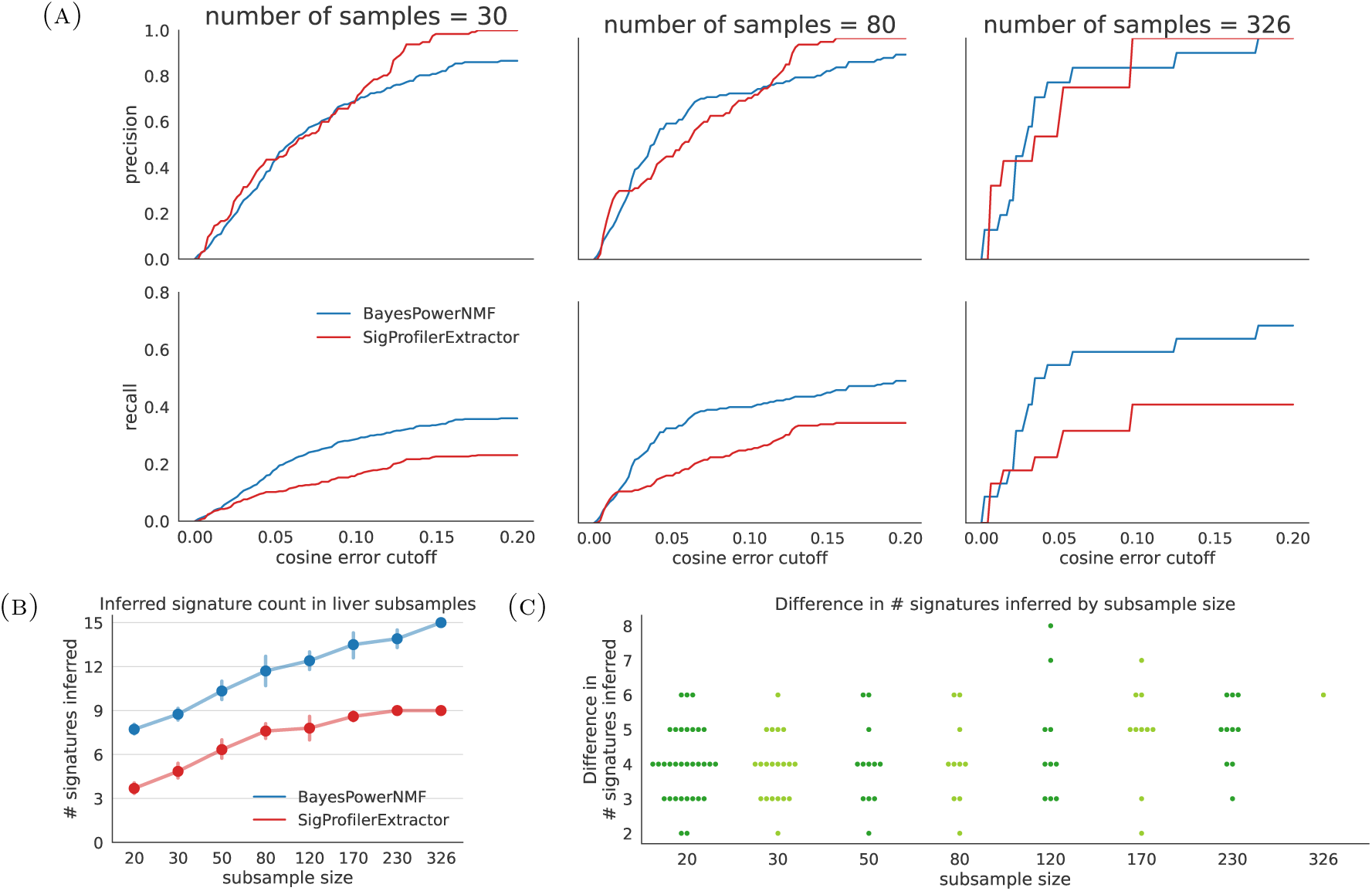
Results for subsampled liver cancer data sets of increasing size. **(A)** Precision and recall of matched signatures, as a function of the cosine error threshold for defining a correct match. BayesPowerNMF almost always strictly dominates SigProfilerExtractor in recall at all sample sizes, and has comparable or better precision for cosine error thresholds values of practical utility (*<* 0.1). **(B)** Number of signatures estimated (with whiskers representing 95% confidence intervals) at each sample size. **(C)**Difference between number of signatures estimated by BayesPowerNMF and SigProfilerExtractor for each subsampled data set. BayesPowerNMF recovers more signatures than SigProfilerExtractor at all sample sizes. The true number of signatures is 21.

To examine the effects of sampling variability, Figure 6c shows the difference in the number of signatures inferred by BayesPowerNMF and SigProfilerExtractor for each individual subsampled data set. On every individual data set, BayesPowerNMF always recovers at least 2 more signatures than SigProfilerExtractor, and sometimes as many as 7 or 8 more.

### 3.4. Summary of simulation results

Our simulation results demonstrate that BayesPowerNMF has improved performance compared to leading methods—enabling the discovery of a greater number of meaningful signatures—while also being robust to plausible model misspecification. One downside of BayesPowerNMF is that the additional robustness obtained in misspecified settings comes at a cost of reduced performance in well-specified settings compared to using the standard Bayesian posterior; see Figure S1. Overall, we find that BayesPowerNMF has the following specific advantages.

#### Higher precision

Compared to SigProfilerExtractor and SignatureAnalyzer, BayesPowerNMF tends to have higher precision. This holds across different DGPs (Figures 3 and S3a) and across sample sizes (Figure 6a), especially for smaller cosine error cutoffs that would produce recognizable signature matches (for instance, 0.05 or 0.1).

#### Higher recall

BayesPowerNMF consistently outperforms the competitors in terms of recall, particularly in misspecified cases (Figures 3 and 6a and Table S1). In the misspecified cases, SigProfilerExtractor tends to be too conservative, inferring too few signatures (Figure 3 and Table S2). SignatureAnalyzer sometimes infers more signatures, but many of the estimated signatures either have very large cosine error (*>* 0.2) or do not even get matched to one of the true signatures used to generate the data.

#### Uncertainty quantification

As a fully Bayesian method, BayesPowerNMF quantifies uncertainty in the signatures as well as the loadings via posterior samples produced by MCMC (Figures 5a and 5b). We find that the model’s reported uncertainty in the signatures is correlated with the actual recovery error (Figure S3b), indicating that the uncertainty quantification is meaningful. In contrast, SigProfilerExtractor or SignatureAnalyzer do not provide uncertainty quantification in either the signatures or the loadings.

#### Few spurious signatures

We consider an estimated signature to be “spurious” if it is matched to a COSMIC signature that was not used to generate the simulated data set.

Out of the 15 complete simulated data sets we consider, BayesPowerNMF infers a spurious signature in just one data set (see Tables S1 and S2). SignatureAnalyzer inferred spurious signatures in several synthetic data sets, even in the well-specified case (see Table S1).

## 4. Application

We compare the results of BayesPowerNMF and SigProfilerExtractor on whole-genome sequencing (WGS) data from the PCAWG project (The ICGC/TCGA Pan-Cancer Analysis of Whole Genomes Consortium, 2020). SignatureAnalyzer is excluded from this comparison due to its overall poor performance. We consider the PCAWG WGS data for the same six cancer types as in the simulation study: lung adenocarcinoma (*N* = 38), stomach (*N* = 75), melanoma (*N* = 107), ovary (*N* = 113), combined breast cancers (*N* = 214), and liver (*N* = 326). While any set of mutation types could be used to define the input count data matrices, we consider the 96 types of single-base substitutions (SBSs) since these are the most commonly used set for mutational signatures analyses. We run BayesPowerNMF and SigProfilerExtractor on the *N* × 96 count data matrix for each cancer type separately.

To evaluate performance, we use the COSMIC v3 signatures^2^ as a proxy for the true signatures. (Note, however, that we still use COSMIC v2 in the BayesPowerNMF workflo when defining pilot signatures for calibration.) The COSMIC signature database is a curated reference set of mutational signatures that is commonly used in cancer genomics studies. The current versions of the COSMIC signatures are based on 2,780 cancer genomes from the PCAWG project—along with a number of other available cancer genomes—spanning a wide range of cancer types (Alexandrov et al., 2020; The ICGC/TCGA Pan-Cancer Analysis of Whole Genomes Consortium, 2020). The COSMIC v3 signatures were defined by running a previous iteration of SigProfilerExtractor on each cancer type separately, and combining the results to construct a set of consensus signatures. Thus, one would expect SigProfilerExtractor to perform very well in these comparisons, since it was instrumental in defining our proxy for ground truth.

Since the actual ground truth is unknown on real data, we quantify method accuracy using using three proxies for quality. First, we compare the number of signatures inferred by each method on each cancer type. For a given amount of data, it is desirable to infer more signatures since this indicates improved recall. However, it is also important that the inferred signatures correspond to true signatures and are accurately estimated, in other words, it is important to have high precision. Thus, second, we compute the error with which the COSMIC signatures are reconstructed. More precisely, we assign the inferred signatures to the best-matching COSMIC v3 signatures using the Hungarian algorithm, and compute the cosine error between each matched pair. It is desirable to have smaller error, since this indicates greater accuracy in recovering the current best understanding of ground truth. Third, we compare the estimated loadings for each method to the loadings presented by the PCAWG consortium, which they obtained by running non-negative least squares (NNLS) using the complete set of COSMIC signatures, rather than using NMF. We refer to these as the “COSMIC+NNLS” loadings.

### 4.1. Application results

Figure 7 summarizes the results; see the description of Figure 4 and its caption for the interpretation of this plot.

**Figure 7.**
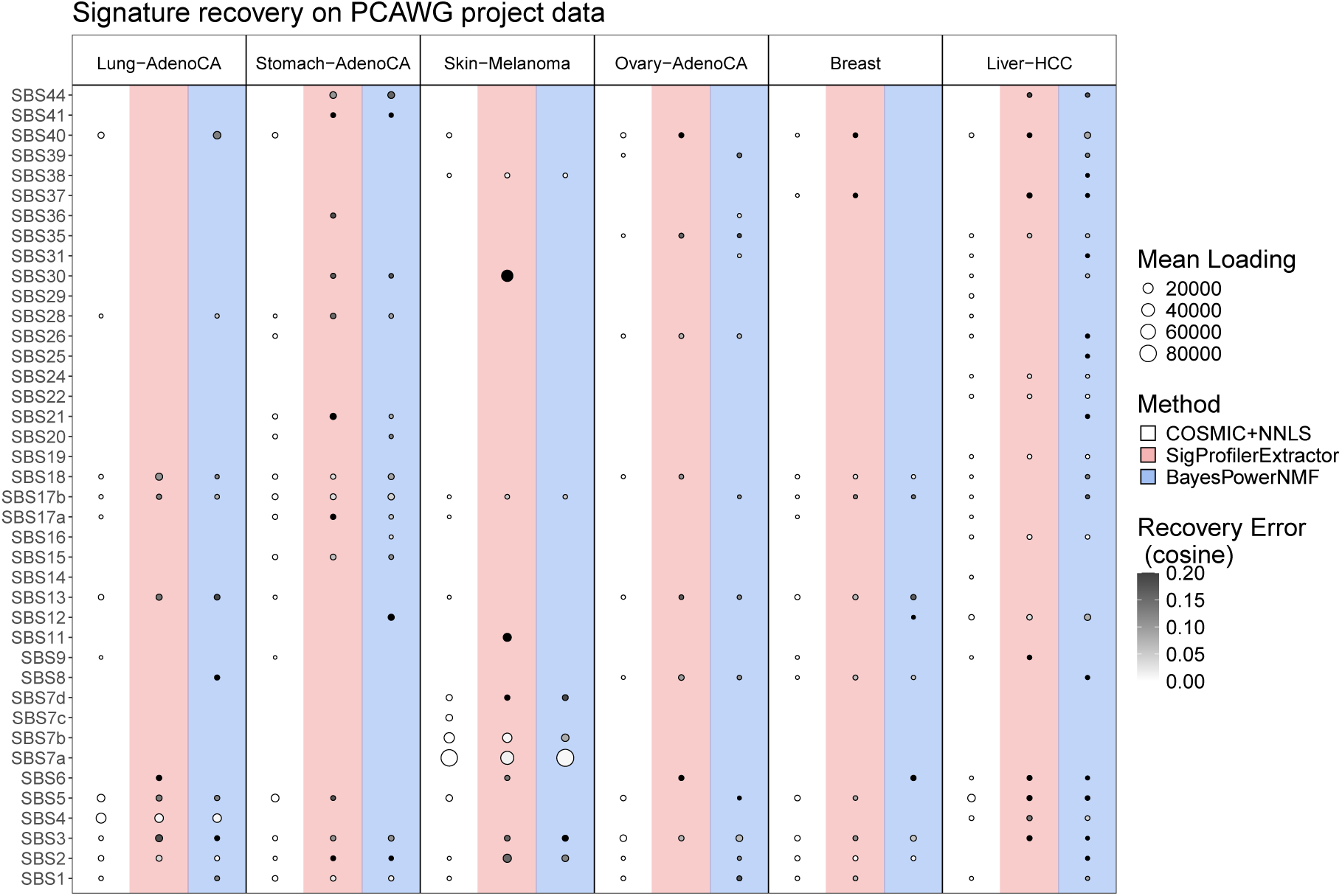
Mean loading and cosine error between each estimated signature and its matching COSMIC v3 signature. Results are shown for each method applied to the PCAWG data for each of six cancer types. See the description of Figure 4 for explanation of the plot. COSMIC+NNLS uses non-negative least squares to estimate loadings for the COSMIC v3 signatures.

#### Number of signatures inferred

In four of the six cancer types, BayesPowerNMF estimates more signatures than SigProfilerExtractor (Figure 7 and Table S3). Furthermore, in one of the two cancer types where SigProfilerExtractor infers more signatures (melanoma), some of the estimated signatures appear to be near duplicates rather than authentically different signatures (Figure S5). In many cases, BayesPowerNMF recovers almost as many signatures as COSMIC+NNLS (Figure 7 and Table S3), even though BayesPowerNMF is only using a small subset of data from one cancer type whereas COSMIC+NNLS is effectively using ≈ 3,000 samples.

#### Reconstruction error

Both methods perform roughly equally well in terms of the accuracy with which they reconstruct COSMIC signatures. Indeed, the difference between the average cosine errors for BayesPowerNMF and SigProfilerExtractor (0.146 and 0.133, respectively) is not statistically significant (*p* = 0.9265, Mann–Whitney rank test). This is despite the fact that (1) SigProfilerExtractor is at an advantage since the COSMIC signatures were constructed using its algorithm and (2) BayesPowerNMF recovers a greater proportion of signatures (Table S3).

#### Loadings

In most cases, the loadings estimated by BayesPowerNMF and SigProfilerExtractor are of similar magnitude to one another and to the COSMIC+NNLS loadings, at least for the signatures that they have in common (Figure 7). However, on the melanoma data, SigProfilerExtractor gives large loadings to two apparently spurious signatures that have high cosine error with their nearest COSMIC v3 matches (SBS30 and SBS11).

Overall, the real data results lead to similar conclusions as the simulation study results. These findings indicate that our proposed method provides improved accuracy and robustness to model misspecification compared to the state-of-the-art, both on synthetic examples and real biological data. It is notable that SigProfilerExtractor appears to perform more poorly on the real PCAWG data than on our simulated misspecified counts data, compared to BayesPowerNMF. This suggests that the modes of misspecification in real mutation counts data are more complex than is captured in each of our DGPs for misspecified counts, but this is accounted for in BayesPowerNMF due to the flexibility of the power posterior.

Finally, we note that the BayesPowerNMF workflow uses COSMIC v2 for the pilot signatures even on this PCAWG data, where we ultimately compare to COSMIC v3. BayesPowerNMF still successfully infers v3 signatures on this data, including ones with no corresponding signature in v2 (specifically, SBS31 and above in Figure 7). This suggests that the precise choice of pilot signatures does not strongly affect the final results from BayesPowerNMF.

## 5. Discussion

Our results elucidate specific shortcomings of two of the standard methods for mutational signature discovery, SigProfilerExtractor and SignatureAnalyzer, and highlight the benefits of carrying out thorough simulation studies to better understand the failure modes of any given method before proceeding to analysis of real data. SigProfilerExtractor uses a consensus bootstrap approach, which may explain why it tends to miss more signatures with small loadings (Figure S2). Specifically, since each bootstrap sample uses only ≈ 63% of the data on average, it makes sense that SigProfilerExtractor would have lower power to discover mutational processes, particularly when they have smaller loading (Figure S2). BayesPowerNMF does not have this issue since it uses all of the available data. This suggests that the SigProfilerExtractor results might be improved by using a version of bootstrap with continuous weights, such as the Bayesian bootstrap, rather than multinomial bootstrap.

Furthermore, while SigProfilerExtractor is generally more conservative in the sense that it recovers fewer signatures than BayesPowerNMF, it can still overfit the data in cancer types with high mutational burden (such as melanoma, see Figure S5) or many samples (such as liver). This makes sense, since these are the cases in which we would expect the negative effects of misspecification to be most pronounced (Miller and Dunson, 2018). BayesPowerNMF avoids this issue by employing a power posterior to improve robustness to misspecification.

Similarly, a disadvantage of SignatureAnalyzer is that it relies heavily on the correctness of the Poisson NMF model (Zito and Miller, 2024). While SigProfiler’s use of bootstrapping can somewhat mitigate the effects of misspecification (Huggins and Miller, 2023, 2024) and BayesPowerNMF’s use of the power posterior provides robustness, SignatureAnalyzer has no mechanism to compensate for model misspecification. As a result, it appears that SignatureAnalyzer is particularly negatively affected by misspecification. Furthermore, SignatureAnalyzer uses an estimation algorithm based on several heuristic approximations to the objective function (Tan and Févotte, 2013). The use of these heuristics may explain why SignatureAnalyzer was not able to accurately recover the true signatures even on the simulated well-specified data sets, a setting in which we would expect a model-based method to perform well. In contrast, BayesPowerNMF does not suffer from this issue, since it does not employ any approximations other than MCMC sampling. Consequently, BayesPowerNMF exhibits good performance on the well-specified data, as expected.

The main disadvantage of BayesPowerNMF is that, with currently available inference techniques, it becomes computationally prohibitive for larger data sets. Improving the scalability of BayesPowerNMF is an important direction for future work. A related disadvantage is the somewhat complicated workflow for selecting the power *ξ* to be used in the power posterior. It would be preferable to have a simpler method of determining an appropriate power for a given set of plausible data generating processes.

Overall, BayesPowerNMF yields superior performance by *(i)* using all the available data and employing a full Bayesian model to extract as much information as possible from the data, while *(ii)* using a power posterior to obtain robustness to model misspecification without assuming a particular form of misspecification. More generally, our methodology can be used to perform non-negative matrix factorization for other applications as well, since it is not limited to mutational signatures analysis. It would be interesting to explore other applications of the method in future work.

## Acknowledgments

C.X. was supported by NIH Training Grant T32GM135117 and NSF Graduate Research Fellowship DGE-2140743. J.W.M. and S.L.C were supported in part by the National Cancer Institute of the National Institutes of Health under award number R01CA240299. J.H.H. was supported by the National Institute of General Medical Sciences of the National Institutes of Health under award number R01GM144963 as part of the Joint NSF/NIGMS Mathematical Biology Program. The content is solely the responsibility of the authors and does not necessarily represent the official views of the National Institutes of Health.

## Appendix A. Additional empirical results

**Figure S1.**
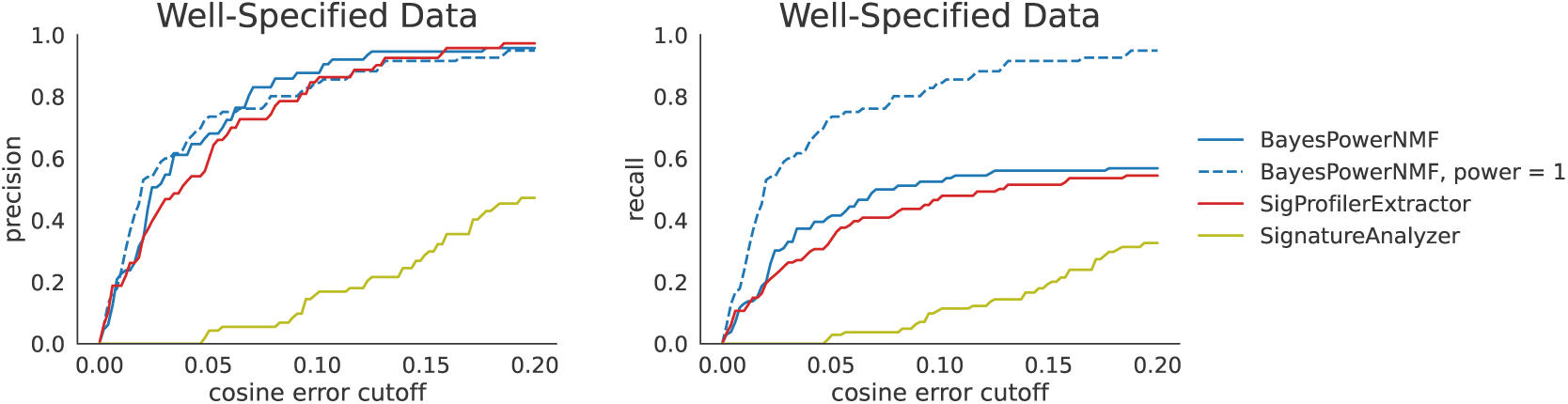
Same as Figure 3 but also showing the precision and recall for the standard Bayesian posterior (blue dashed lines) for the same NMF model. This corresponds to BayesPowerNMF with a power of *ξ* = 1. When the model is correct (that is, in the well-specified case), the standard posterior exhibits higher recall than the power posterior, while maintaining comparable precision.

**Figure S2.**
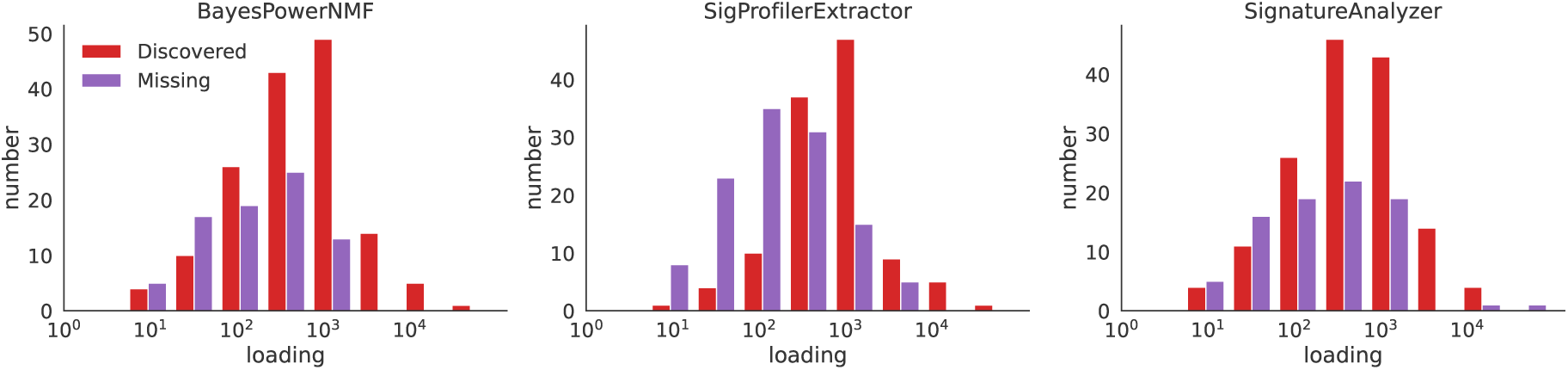
Histograms of ground-truth loadings for pilot signatures discovered and missed by BayesPowerNMF, SigProfilerExtractor, and SignatureAnalyzer across all simulated data sets. SigProfilerExtractor systematically misses many more signatures with small mean loading in the pilot data set.

**Figure S3.**
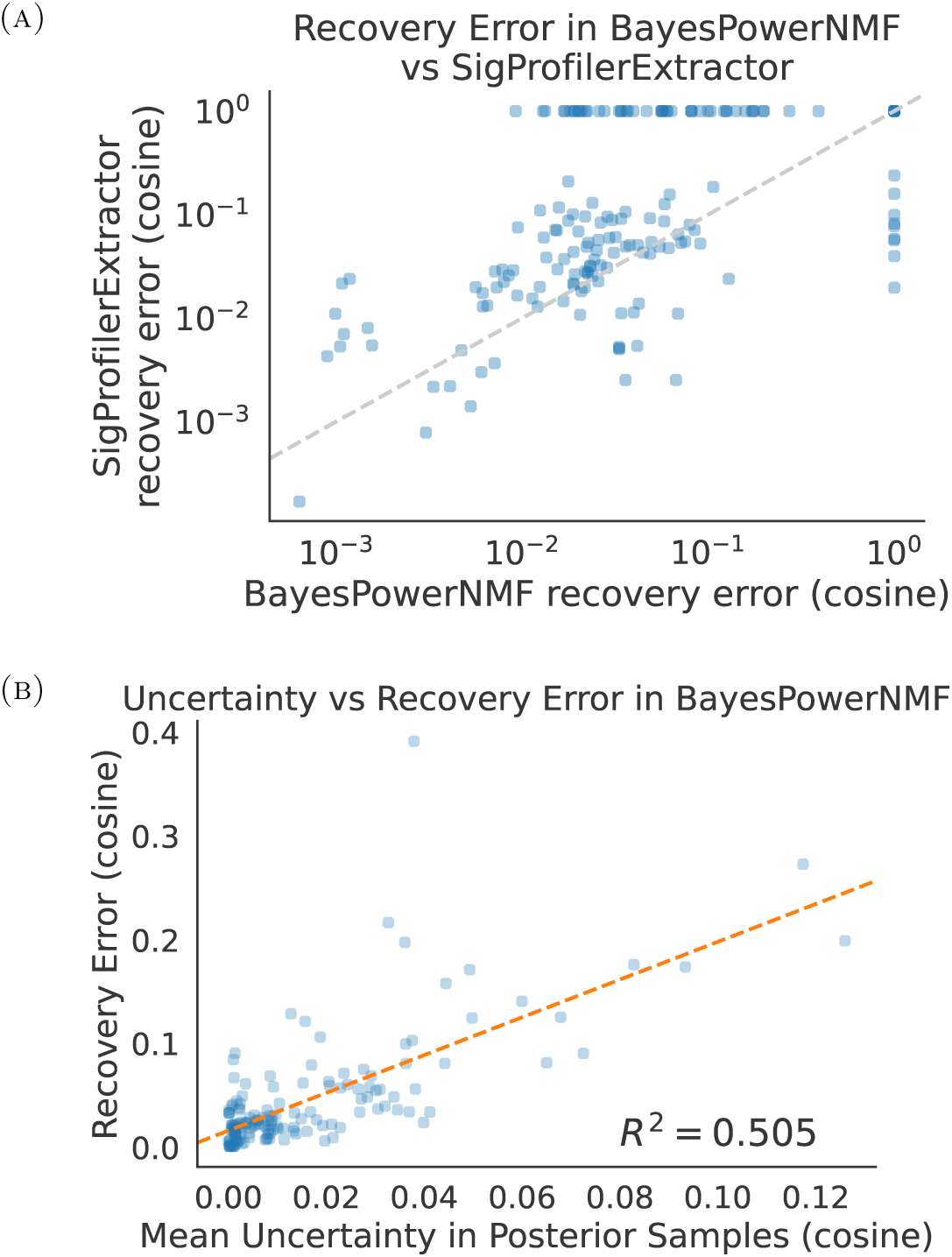
**(A)** Recovery error for SigProfilerExtractor versus BayesPowerNMF. For each of the 15 complete synthetic data sets (see Section 3), for each ground truth signature, we plot the cosine error between the true signature and the signatures estimated by SigProfilerExtractor (y-axis) and BayesPowerNMF (x-axis); a cosine error of 1 indicates that no matching signature was inferred by that method. SigProfilerExtractor misses many of the signatures recovered by BayesPowerNMF (see dots along the top), and has higher cosine error for most of the signatures recovered by both methods (see dots above the diagonal). **(B)** In simulations with BayesPowerNMF, the posterior uncertainty in each signature is correlated with the cosine error between the estimated signature and the ground truth signature. This indicates that the uncertainty quantification is providing meaningful information about the actual error. Here, uncertainty is defined as the mean cosine error between posterior samples and the posterior mean, for each signature.

**Figure S4.**
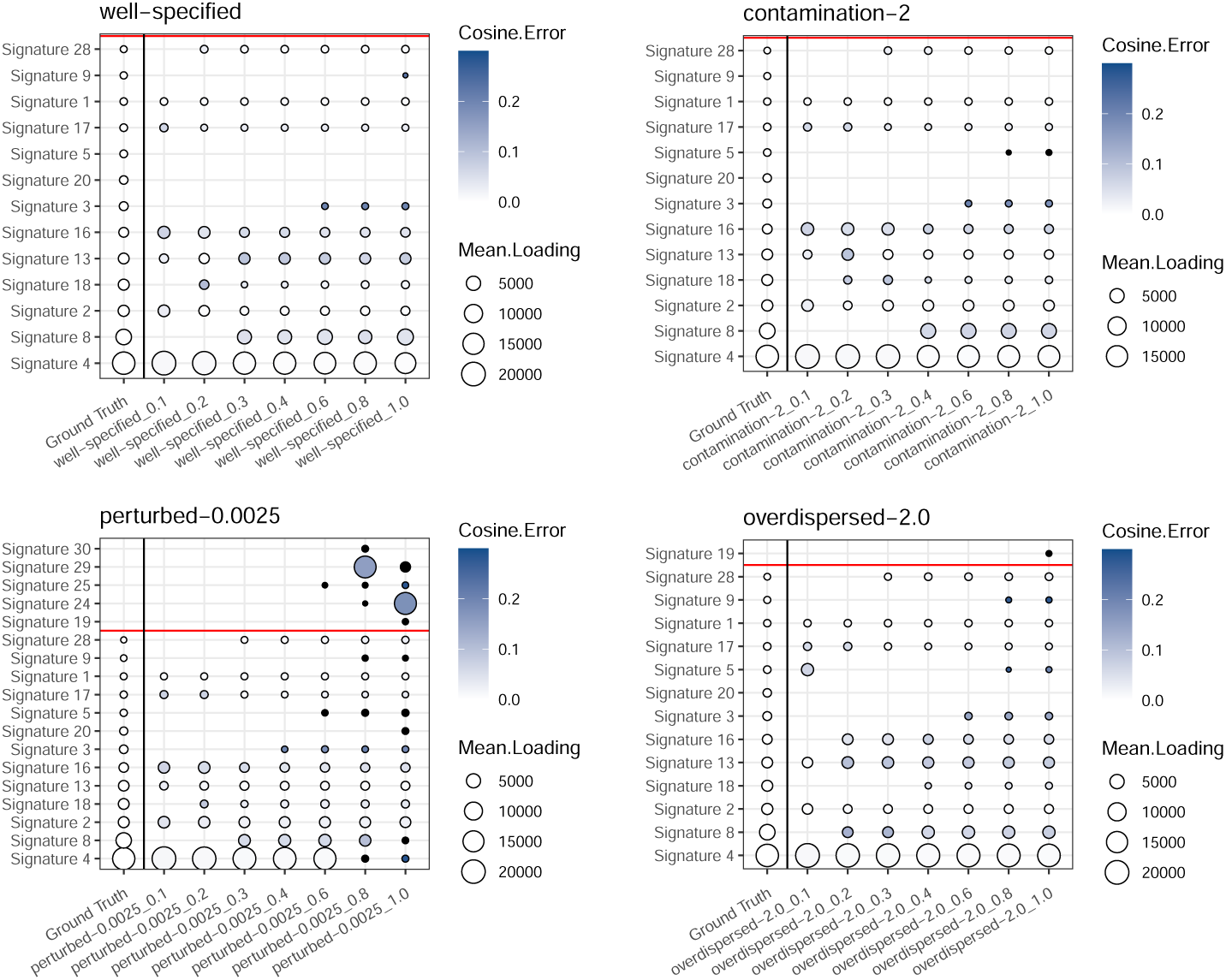
Bubble plots used to select the power *ξ* in stage 4 of the BayesPowerNMF workflow. Each panel corresponds to one simulated data set. See the description of Figure 4 for the interpretion of this type of plot.

**Figure S5.**
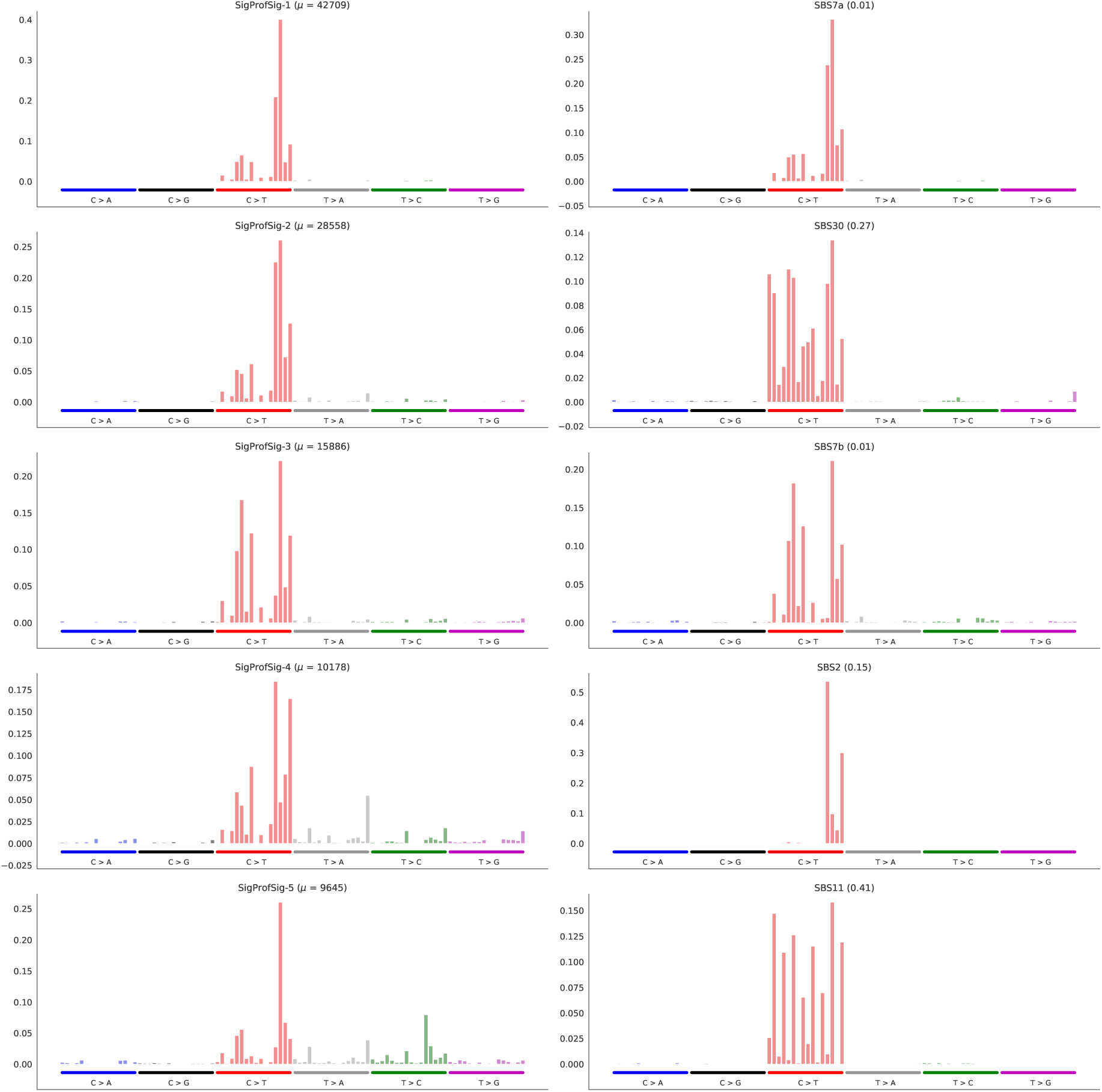
The top 5 out of 10 signatures inferred by SigProfilerExtractor from melanoma mutation counts from the PCAWG project (left) and their best match reference signatures from COSMIC v3 (right). There appears to be potential duplication between (a) SigProfSig-1, SigProfSig-2, and SigProfSig-5, and (b) SigProfSig-3 and SigProfSig-4, in the sense that these may be slightly perturbed versions of the same “true” signature.

**Figure S6.**
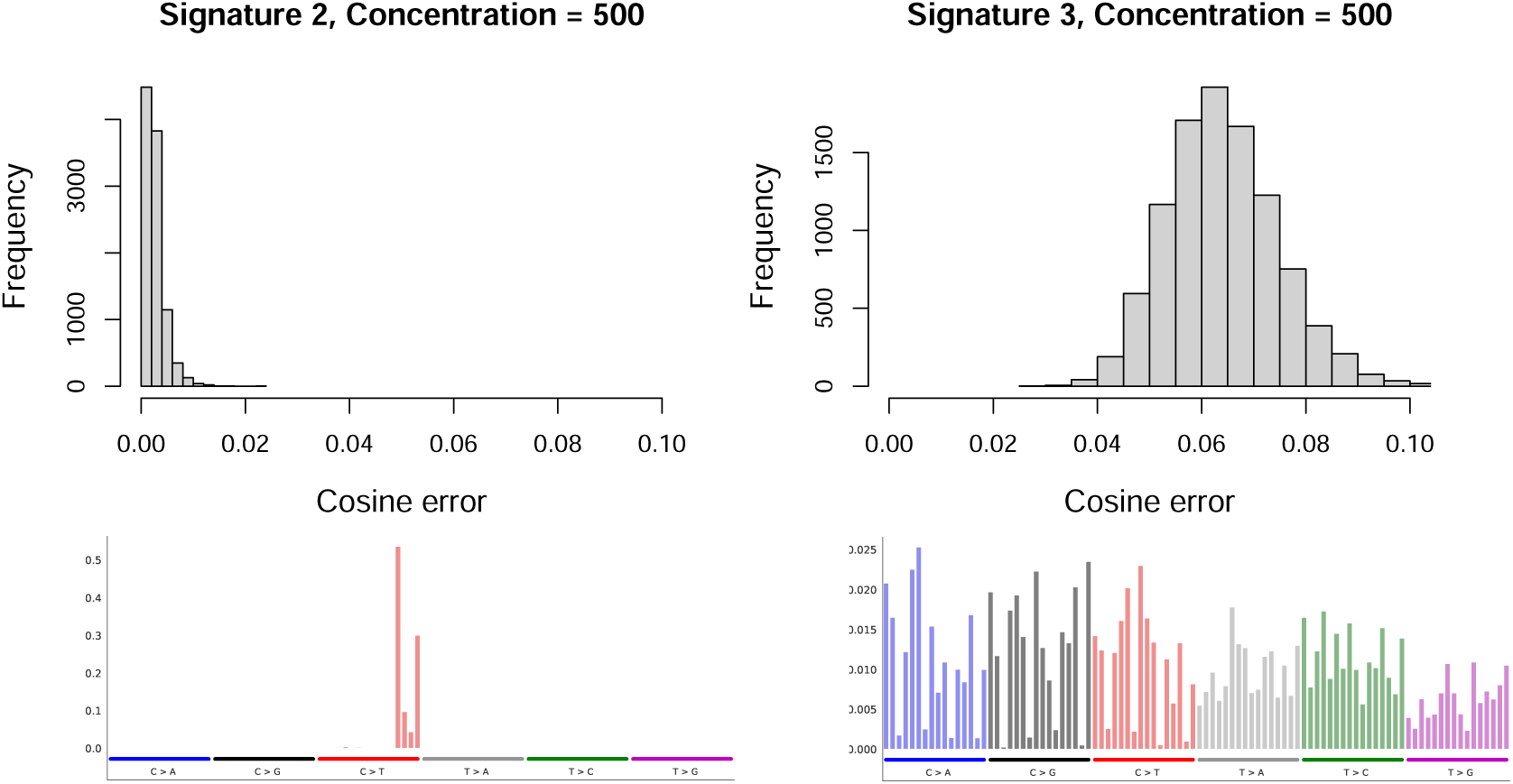
(Top) Distribution of cosine errors between *R*^*^ and 10,000 samples from Dirichlet(*βR*^*^), where *β* = 500 and *R*^*^ is Signature 2 (left) or Signature 3 (right) from COSMIC v2. (Bottom) Mutation frequency profile for Signature 2 (left) and Signature 3 (right).

**Table S1.** Mutational signature discovery results for simulated data, comparing SigProfilerExtractor, SignatureAnalyzer, and BayesPowerNMF on well-specified data for six cancer types. The number of COSMIC v2 signatures used to generate the data is in parentheses after each cancer type. In each entry of the form X +Y *+Z* in the table, X = the number of estimated signatures that were matched to a ground truth signature with a cosine error of *<* 0.2, Y = the number of estimated signatures that were matched to a ground truth signature with a cosine error of ≥ 0.2, and *Z* = the number of estimated signatures that were matched to a COSMIC v2 signature that was not used to generate the data. Bold indicates the best performing method in each case. (SPE = SigProfilerExtractor; SA = SignatureAnalyzer; BPN = BayesPowerNMF; BPN_ξ=1_ = BayesPowerNMF with power of 1, which corresponds to the standard posterior.)

APPENDIX A. ADDITIONAL EMPIRICAL RESULTS
|  | Lung (13) | Stomach (21) | Skin (14) | Ovary (12) | Breast (12) | Liver (21) |
| --- | --- | --- | --- | --- | --- | --- |
| SPE | 7 | 11 | 6 | 7 | **10** | 9 |
| SA | 5 +4 | 5 +8 +1 | 6 +4 +4 | 5 +2 | 3 +5 +2 | 8 +6 |
| BPN | **9** | 10 +1 | 5 +1 | 7 | 7 | 15 |
| BPN<sub>$\xi=1$</sub> | 9 +2 | **14** +1 | **13** | **10** | 12 +3 | **16** +1 |

**Table S2.** Same as Table S1 but for both the well-specified and misspecified settings, for three cancer types.

|  | Lung (13) |  |  | Ovary (12) |  |  | Liver (21) |  |  |
| --- | --- | --- | --- | --- | --- | --- | --- | --- | --- |
|  | SPE | SA | BPN | SPE | SA | BPN | SPE | SA | BPN |
| well-specified | 7 | 5 +4 | **9** | **7** | 5 +2 | **7** | 9 | 8 +6 | **15** |
| α-contaminated | 7 | 5 +4 | **9** | 5 +1 | 5 +2 | **5** | 9 | 7 +5 | **15** |
| γ-perturbed | 7 | 3 +7 +1 | **10** | 5 +1 | 5 +2 | **8** | 9 | 10 +5 | **17** |
| κ-overdispersed | 7 | 4 +5 | **9** | 5 | 5 +2 | **9** +1 | 9 | 8 +7 | **15** |

**Table S3.**
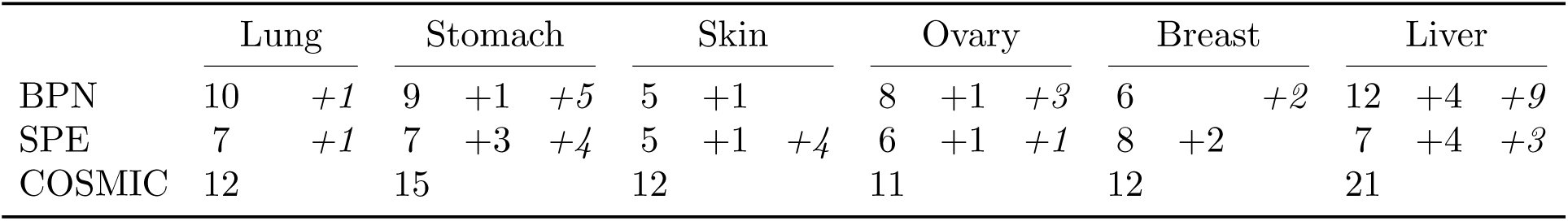
Number of signatures inferred by each method on the PCAWG data for six cancer types, treating COSMIC+NNLS as ground truth. (SPE = SigProfilerExtractor; BPN = BayesPowerNMF; COSMIC = COS-MIC+NNLS.)

## Appendix B. Details on *γ*-perturbed data-generation process

Näıvely, one might try to simulate perturbed signatures by sampling from Dirichlet(*βR*^*^) using the same concentration parameter *β >* 0 for all pilot signatures *R*^*^. However, in terms of cosine error, this leads to very different magnitudes of perturbation depending on the sparsity or flatness of *R*^*^; see Figure S6.

We quantify the “flatness” of a signature *R*^*^ via

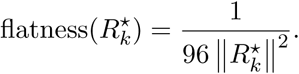

Larger values indicate a flatter signature and smaller values indicate a spikier signature. This expression is specifically inspired by the form of the cosine error formula, and takes values in [0, 1]. Through simulations using each *R*^*^ in the set of COSMIC v2 signatures, we observe that for a range of concentration parameters *β*ranging from 500 to 10,000, the mean cosine error for Dirichlet(*βR*^*^) has a very strong positive linear correlation with flatness(*R*^*^) (Figure S7a).

Further, the slopes of the best-fit line obtained with ordinary least squares (OLS) are strongly linearly correlated with *β* on a log-log scale (Figure S7b).

Based on these observations, we use OLS to fit the empirical relationship between flatness(*R*^*^), the desired mean cosine error *γ* ∈ (0, 1), and the concentration parameter *β*, specifically,

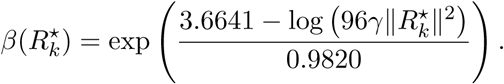

**Figure S7.**
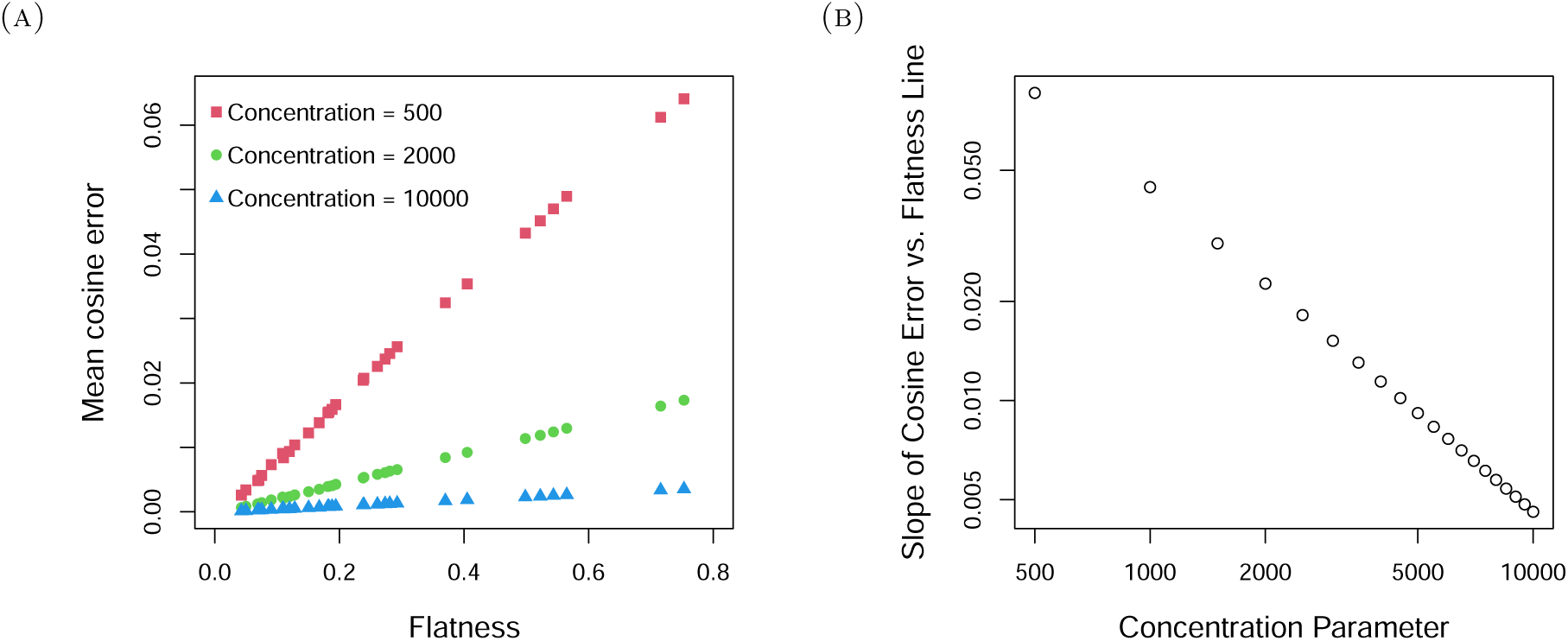
**(A)** Strong linear relationship between flatness(*R*^*^) and the mean cosine error between a sample from Dirichlet(*βR*^*^) and *R*^*^ itself for several different values of *β*. **(B)** Strong linear relationship between the slope of the lines in Figure S7a and the concentration parameter *β* on a log scale.

This relationship allows us to choose a signature-specific concentration parameter to simulate new signatures, while controlling the resulting mean cosine errors across all signatures. We use the function *β*(*R*^*^) to set the values of *β*_k_ for the *γ*-perturbed DGP in Section 2.4.

## Footnotes

1 https://cancer.sanger.ac.uk/signatures/signatures_v2/

2 https://cancer.sanger.ac.uk/signatures/downloads/

